# D-cycloserine (DCS) is Not Susceptible to Self-administration, unlike S-ketamine Using an Intravenous Self-administration Model in Naive and Ketamine-habituated Sprague-Dawley Rats

**DOI:** 10.1101/2022.08.12.503713

**Authors:** Daniel C. Javitt, Jonathan C. Javitt

## Abstract

**OBJECTIVE:** N-methyl D-aspartate Receptor (NMDAR) antagonist antidepressants have known potential for abuse liability. The aim of this study was to evaluate the abuse liability of D-cycloserine (DCS), using a self-administration paradigm in which DCS was tested in its efficacy in substituting for ketamine in ketamine-dependent rats.

**METHODS:** A standard Intravenous self-administration study was conducted in Male adult Sprague-Dawley rats. model to study compounds’ abuse liability. Potential for self-administration was assessed in ketamine-habituated subjects. Subjects were trained to press a lever to obtain food, prior to connection of the lever to intravenous drug administration apparatus. DCS was provided for self-infusion by test subjects at doses of 1.5, 5.0, and 15mg/kg per lever press.

**RESULTS:** S-Ketamine was seen to substitute for ketamine and to result in self-administration at the same frequency. DCS was not seen to result in any self-administration at any of the test doses. The self-infusion behavior of DCS was the same as that of saline.

**CONCLUSION:** C-cycloserine, an a mixed agonist/antagonist of the NMDAR glycine site, which has been shown to have antidepressant and anti-suicidal properties in clinical studies has no apparent potential for abuse liability in a standard rodent self-administration model.

## 1 INTRODUCTION

Functional antagonists of the N-methyl D-aspartate Receptor (NMDAR) were first hypothesized to have antidepressant effects by Trullas and Skolnick.^1^ The discovery that ketamine has a rapid and profound effect on depression and suicidality^2^ has led to broad recognition that glutamate system and the antagonists may also play an important role in depression and suicidality. Ketamine has since been shown in multiple randomized clinical trials to induce nearly immediate remission from depressive symptoms as well as from suicidal ideation.

Despite the potential clinical value of NMDAR-targeted antidepressants, ketamine^3,4^, Dextromethorphan^5^ and other NMDAR antagonist agents have demonstrated potential for abuse and addiction. Recently, the S-enantiomer of ketamine has been demonstrated to have greater potential for self-administration and abuse than the R-enantiomer.^6^ The American Psychiatric Association and other expert bodies have expressed concern about the repeated use of ketamine for the treatment of depression and/or suicidality. The US Food and Drug Administration has issued guidance requiring rodent behavioral studies prior to human trials of any investigational drug known to have CNS activity^7^

D-cycloserine (DCS) is a broad-spectrum antibiotic approved for the treatment of tuberculosis that has been used in millions of individuals without report of significant safety concerns or abuse potential. DCS was serendipitously found to have antidepressant effects by Crane,^8^ which were subsequently confirmed in a placebo-controlled trial.^9^

Following the discovery by Javitt and Zukin of the NMDAR role in phencyclidine-induced psychosis,^10^ the antidepressant effect of DCS was subsequently attributed to it partial agonist effect at the glycine site of the NMDA receptor.^11^ DCS was shown to have potential antidepressant effects on rodent forced swim test by Lopes and coworkers.^12^ DCS at a target dose of >500mg/d has been shown to have a clinical antidepressant effect when administered to patients with treatment resistant depression added to SSRI antidepressants, without hallucinogenic side effects.^13^ DCS sustained the antidepressant effect of ketamine in an open label trial in patients with bipolar depression who were resistant to currently approved pharmacological treatments.^14^ DCS has further been demonstrated to maintain remission from suicidality after ketamine infusion, an effect that is more pronounced among patients with bipolar depression.^15^

DCS is a potentially attractive oral antidepressant candidate because unlike direct NMDA channel blocking agents, it has shown no clinical potential for neurotoxicity or addiction. In this study, we demonstrate that DCS appears to have no potential for abuse liability or self-administration in a standard nonclinical model.

## 2 MATERIAL AND METHODS

All research was performed at PsychoGenics, Inc. (Tarrytown, NY), a facility accredited by the Association for Assessment and Accreditation of Laboratory Animal Care (AAALAC). Procedures were approved by the Institutional Animal Care and Use Committee in accordance with the National Institute of Health Guide for the Care and Use of Laboratory Animals.

### 2.1 Experimental subjects

Male adult Sprague-Dawley rats (300-325 g at arrival) from Envigo Laboratory (Indiana, USA) were used in this study. Upon arrival, the rats were assigned a unique identification number (tail marks). Animals were housed 1 per cage and acclimated for up to 7 days prior to study. All rats were examined, handled, and weighed prior to initiation of the study to assure adequate health and suitability. During the course of the study, 12 h/12 h light/dark cycles were maintained. The room temperature was 20-23°C with a relative humidity maintained 30-70%. Water was provided *ad libitum* for the duration of the study. Following the start of food restriction prior to food training, rats were single housed and food-restricted after recovery with their body weight maintained at 85% of their free-feeding age-matched control body weight.

### 2.2 Test Compounds

Ketamine hydrochloride and S-Ketamine (1.82 mg/ml, equivalent to 0.5 mg/kg/infusion under 350 g body weight and 0.1 ml/infusion rate) was dissolved in saline. The resulting solution was clear. DCS (1.5, 5.0, and 15.0 mg/kg/i.v.infusion) was dissolved in saline. The resulting solutions were clear. Saline was used as vehicle. Ketamine, S-ketamine, or DCS solutions were intravenously self-administered during the 1hr test sessions at a rate of 0.1ml/infusion.

### 2.3 Intravenous Self-administration

#### Apparatus

Intravenous drug self-administration and food-maintained responding training and test took place in operant chambers within sound-attenuating cubicles equipped with an exhaust fan (Med Associates, VT). Each chamber contained two levers situated on one wall of the chamber. Only one of the two levers was active (located on the left side). Pressing the active lever twice caused delivery of reinforcer (in this case ketamine in self administration study). The other lever was “inactive”, i.e. pressing it did not deliver any reinforcement. A stimulus light was located above each lever, but only the one above the active lever was on during the timeout period (defined below). A house light (providing illumination) was located at the top of opposite wall. An infusion pump mounted above each chamber delivered drug solution via Tygon tubing connected to a single channel fluid swivel, which was mounted on a balance arm above the operant chamber. The output of the liquid swivel was attached to the externalized terminus of the intravenous catheter.

#### Food Training and Surgery

Prior to intravenous catheterization, animals were trained to press the active lever to obtain food. Food training started after the rats were food-restricted and reached 85% of the free-feeding body weight after about one week. After acquiring the lever-press response to obtain food through about one week training, rats were implanted with intravenous catheters. Catheters were flushed with 0.2 ml of heparin-gentamicin solution every day to avoid clogging and ensure smooth drug infusion. The rats were on free feeding two days prior to surgery and throughout recovery,

#### Ketamine Acquisition

Ketamine acquisition, i.e. generation of ketamine dependence was the first step of self-administration in this study. One week after the surgery, single housed rats were food restricted again and maintained at 85% of their free-feeding age-matched control body weight throughout the study. After rats reached 85% of the body weight, rats were allowed to self-administer ketamine solution by pressing the active lever in a fixed-ratio (FR) schedule of reinforcement. In this study we used FR2, i.e. two lever presses for one ketamine delivery. The dose of ketamine was 0.5 mg/kg/infusion. Each ketamine infusion lasted 1.0 sec. Delivery of ketamine was followed by a 20-second timeout period, during which no drug was delivered even if the active lever was pressed. During timeout, the stimulus light above the active lever was on.

#### Ketamine Substitution

After ketamine self-administration was achieved through 17 days of training, the rats experience a 4-day extinction period (saline infusion only), and then the rats were tested with S-ketamine 0.5mg/kg/infusion and different doses of DCS (1.5, 5.0, and 15.0 mg/kg/infusion). One dose was tested each week. In substitution weeks, rats were always trained with ketamine (0.5 mg/kg/infusion, same as acquisition session) on Monday, and then with the S-ketamine or one dose of DCS on Tuesday to Fridays. The test session lasted 1 hour with FR2 drug delivery schedule as described above in the ketamine training session.

During the study procedure, Methohexital sodium (Brevital®, Henry Schein Animal Health, USA,) was used for catheter evaluation and thereby for indirect infusion confirmation. Brevital is a short-acting barbiturate that, when infused through the catheter, produces overt signs of sedation within seconds. The Brevital test (0.2 ml of 1% solution) was performed roughly every other week, usually after compound test on Friday to ensure patency of the catheter.

### 2.4 Statistical Analysis

SigmaStat software package Version 12.5 was used for statistical analysis. One-way repeated measure ANOVA, followed by Fisher LSD *post hoc* test and student t-test were used in different statistical analyses. An effect was considered significant if P<0.05. Data are represented as the mean and standard error to the mean (s.e.m.).

## 3 RESULTS

### 3.1 Acquisition and extinction

Figure 1 shows the ketamine acquisition curve and saline extinction before substitution test. In the present study the rats were ready for substitution test after 17 days of ketamine acquisition training with FR2. Saline extinction was conducted in between the ketamine training and the first substitution test session. As shown in the graph, the lever response was quickly extinguished if the ketamine-trained rats could only obtain saline infusion.

**Figure 1:**
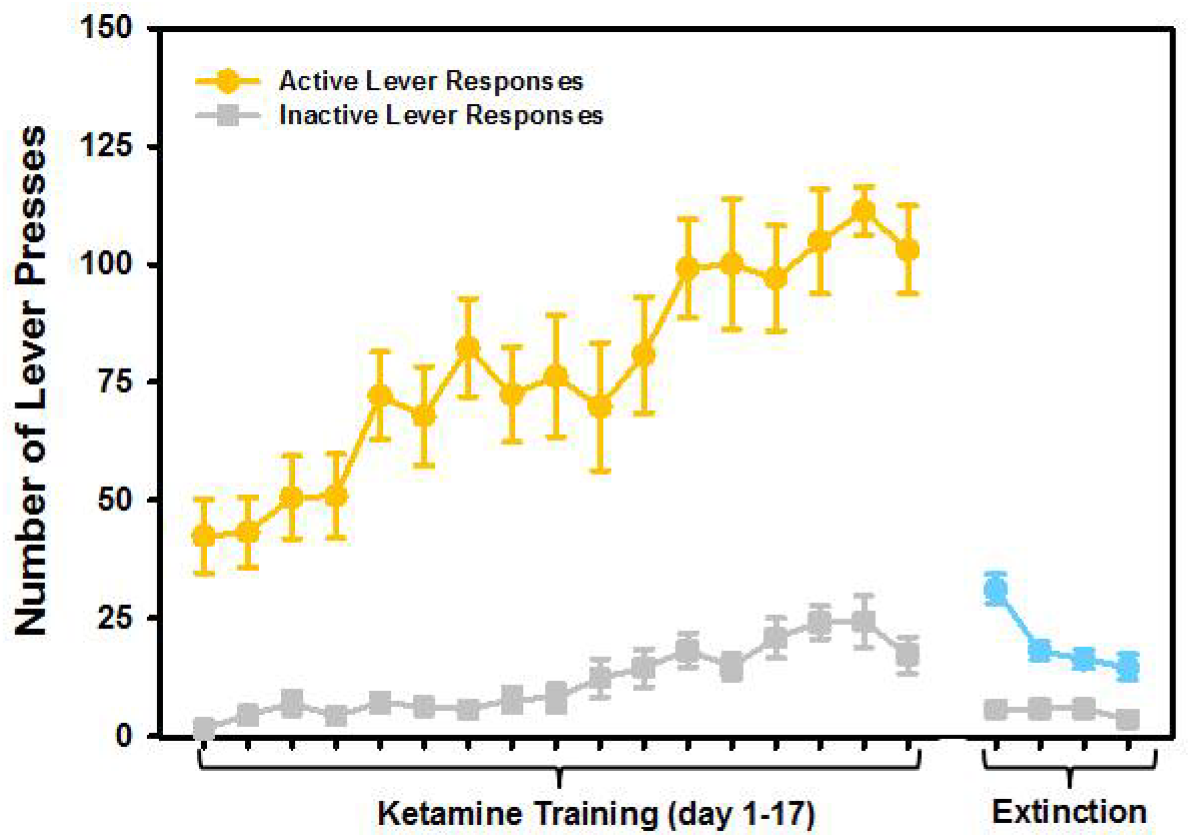
Ketamine (0.5 mg/kg/infusion, FR2) acquisition curve and extinction curve.

Figure 2 shows the average of number of drug infusions (equal to half of the number of positive lever presses) in the last two days of ketamine acquisition, saline extinction, S-ketamine substitution and substitutions of each dose of DCS, respectively. One-way repeated measure ANOVA found a significant treatment effect [F(5,71)=165.162, P<0.001]. *Post hoc* comparison demonstrated that none of the DCS doses showed evidence of substitution for ketamine. The rats’ lever responses in all doses of DCS were only 10% or less of that associated with ketamine infusion and not higher than that under saline infusion. On the other hand, S-ketamine reached the criterion of full substitution (>80%), which indicated the validity of this assay. However, the ~13% less infusion still make S-ketamine treatment significantly different from ketamine treatment (P<0.01), partially due to relatively small deviations in both groups.

**Figure 2:**
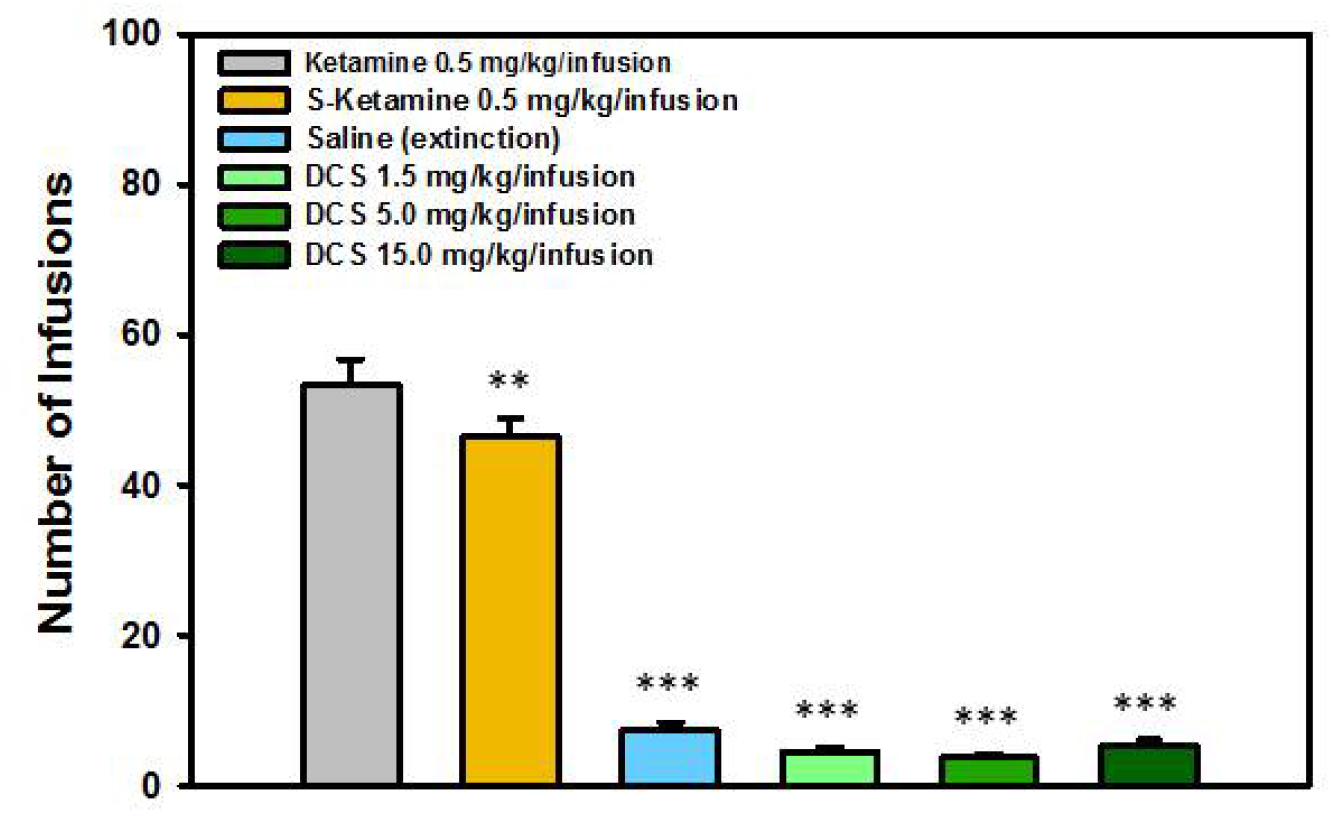
Numbers of infusions in different treatments groups. Data indicated that S-ketamine largely substituted for ketamine, meanwhile DCS did not showed any sign of substitution for ketamine in all doses. (**: P<0.01 and ***: P<0.001 comparing to ketamine treatment).

Figure 3 provides an alternate analysis, which demonstrates the efficacy of S-ketamine and different DCS doses in ketamine substitution. In these analyses we compared ketamine data from the Monday of each week with the corresponding data of DCS and S-ketamine. The data showed that while the ketamine-trained rats were immediately responsive to S-ketamine (P>0.05), their response levels under all DCS doses were significantly lower than that under ketamine (Ps < 0.001).

**Figure 3:**
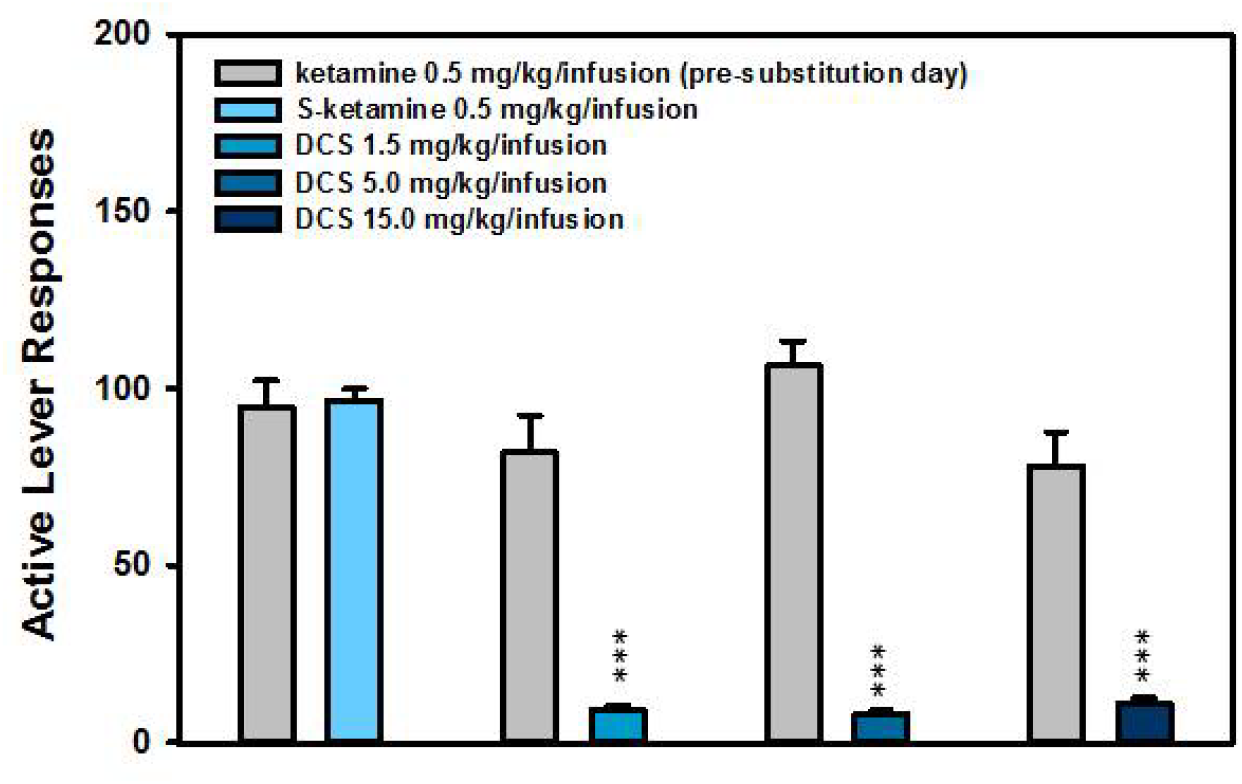
Numbers of Active Lever presses in S-ketamine or different DCS doses relative to corresponding ketamine responses. (***: Ps<0.001, student t-test.)

## CONCLUSION

Current FDA guidance identifies criteria for determining whether human abuse potential must be studied in connection with a CNS-active drug. Those criteria focus on demonstrating whether the investigational drug offers rewarding properties that support animal self-administration and whether abuse-related adverse events (including euphoria-related adverse events) are seen in clinical studies of healthy individuals. ^7^ Although DCS has long been known to cause infrequent cases of hallucination in some patients treated for multi drug-resistant tuberculosis,^16^ those hallucinations have uniformly been viewed as a negative side effect and there are no known reports of DCS-abuse for an hallucinogenic effect. In various studies of DCS, combined with or on top of a 5-HT2A antagonist (either an SSRI antidepressant or atypical antipsychotic), no instance of hallucination has been seen ^13,14,15,17^ and no reports of hallucination have emerged from ongoing clinical trials (NCT03395392).

We evaluated abuse liability of compound D-cycloserine (DCS) using substitution paradigm of intravenous self-administration and determined that DCS does not substitute for ketamine in a self-administration model of ketamine-addicted rodents. Intravenous self-administration is an important animal model to study compounds’ abuse liability. Studies indicated that most drugs that are abused in humans are self-administered by laboratory animals, and most drugs that are not self-administered by laboratory animals are not abused in humans. The present study was designed to assess whether DCS can substitute for ketamine. S-ketamine, the S(+) enantiomer of ketamine, was used as positive control in this study.

The study showed that reference compound S-ketamine demonstrated full substitution on ketamine-dependent rats using a substitution protocol, indicating the validity of this paradigm. On the other hand, DCS in all doses failed to substitute for ketamine in ketamine-dependent rats; rats showed similar frequencies of lever responses to the frequency under saline infusion.

Although there is no direct comparison between an infusion dose in self-administration and a dose through i.p. administration, we may still roughly relate two dosing routes. If the rats infused DCS at the same frequency as they did on ketamine, they could administer about 30, 100 and 300 mg DCS in 1 hour with the three infusing solutions respectively. At 0.35 – 0.4 kg body weight, this would correspond to 75, 250 and 750 mg/kg doses. If DCS had some capacity of substitution for ketamine, rats should have immediately begun self-administration at the first substitution day due to the momentum from ketamine training, following which self-infusion would be expected to increase. The results showed that the ketamine momentum was not effective in leading to substitution, and the rats’ responses to all doses of DCS were comparable to saline extinction.

Our results showing lack of self-administration of DCS may be relevant to concerns previously raised for the clinical use of (S)-ketamine nasal spray if self-administered by patients during treatment^18^, especially in patients with comorbid substance use disorders^19^. Our findings may also have potential clinical implications for the use of DCS in the treatment of depression in preference to ketamine. Specifically, if current clinical trials indicate that DCS shows antidepressant efficacy, the self-administration findings reported herein predict that DCS may have a more favorable risk/benefit treatment profile than (S)-ketamine, especially in patients with comorbid substance use disorders. Based on the data we conclude that DCS has no abuse liability even using the substitution paradigm in nonclinical subjects.

## 4 SUPPLEMENTARY STATISTICAL TABLES

### 4.1 Comparison of infusions in substitution stage

#### One Way Analysis of Variance

| Group Name | N | Missing | Mean | Std Dev | SEM |
| --- | --- | --- | --- | --- | --- |
| saline | 12 | 0 | 7.500 | 3.254 | 0.939 |
| Ketamine | 12 | 0 | 53.375 | 11.790 | 3.403 |
| S-Ketamine | 12 | 0 | 46.542 | 8.497 | 2.453 |
| DCS 1.5 | 12 | 0 | 4.542 | 1.971 | 0.569 |
| DCS 5.0 | 12 | 0 | 3.792 | 1.573 | 0.454 |
| DCS 15.0 | 12 | 0 | 5.375 | 2.524 | 0.729 |
| Source of Variation | DF | SS | MS | F | P |
| Between Groups | 5 | 32279.406 | 6455.881 | 165.162 | <0.001 |
| Residual | 66 | 2579.813 | 39.088 |  |  |
| Total | 71 | 34859.219 |  |  |  |

All Pairwise Multiple Comparison Procedures (Fisher LSD Method):

Comparisons for factor: **Treatment**

All Pairwise Multiple Comparison Procedures (Fisher LSD Method): Comparisons for factor: **Treatment**
| Comparison | Diff of Means | LSD(alpha=0.050) | P |
| --- | --- | --- | --- |
| Ketamine vs. DCS 5.0 | 49.583 | 5.096 | <0.001 |
| Ketamine vs. DCS 1.5 | 48.833 | 5.096 | <0.001 |
| Ketamine vs. DCS 15.0 | 48.000 | 5.096 | <0.001 |
| Ketamine vs. saline | 45.875 | 5.096 | <0.001 |
| Ketamine vs. S-Ketamine | 6.833 | 5.096 | 0.009 |
| S-Ketamine vs. DCS 5.0 | 42.750 | 5.096 | <0.001 |
| S-Ketamine vs. DCS 1.5 | 42.000 | 5.096 | <0.001 |
| S-Ketamine vs. DCS 15.0 | 41.167 | 5.096 | <0.001 |
| S-Ketamine vs. saline | 39.042 | 5.096 | <0.001 |
| saline vs. DCS 5.0 | 3.708 | 5.096 | 0.151 |
| saline vs. DCS 1.5 | 2.958 | 5.096 | 0.251 |
| saline vs. DCS 15.0 | 2.125 | 5.096 | 0.408 |
| DCS 15.0 vs. DCS 5.0 | 1.583 | 5.096 | 0.537 |
| DCS 15.0 vs. DCS 1.5 | 0.833 | 5.096 | 0.745 |
| DCS 1.5 vs. DCS 5.0 | 0.750 | 5.096 | 0.770 |

### 4.2 Comparison of lever responses of ketamine with S-ketamine and DCS

**t-test**

4.2 Comparison of lever responses of ketamine with S-ketamine and DCS: t-test
| Group Name | N | Missing | Mean | Std Dev | SEM |
| --- | --- | --- | --- | --- | --- |
| Ketamine | 12 | 0 | 94.667 | 25.949 | 7.491 |
| S-Ketamine | 12 | 0 | 96.708 | 10.943 | 3.159 |

Difference −2.042

t = −0.251 with 22 degrees of freedom.

95 percent two-tailed confidence interval for difference of means: −18.901 to 14.818

Two-tailed P-value = 0.804

**t-test**

| Group Name | N | Missing | Mean | Std Dev | SEM |
| --- | --- | --- | --- | --- | --- |
| Ketamine | 12 | 0 | 81.833 | 36.581 | 10.560 |
| DCS 1.5 | 12 | 0 | 9.375 | 3.868 | 1.117 |

Difference 72.458

t = 6.824 with 22 degrees of freedom.

95 percent two-tailed confidence interval for difference of means: 50.436 to 94.480

Two-tailed P-value = 0.000000744

**t-test**

| Group Name | N | Missing | Mean | Std Dev |
| --- | --- | --- | --- | --- |
| SEM |  |  |  |  |
| Ketamine | 12 | 0 | 106.417 | 24.322 |
| 7.021 |  |  |  |  |
| DCS 5.0 | 12 | 0 | 8.042 | 3.137 |
| 0.905 |  |  |  |  |

Difference 98.375

t = 13.896 with 22 degrees of freedom.

95 percent two-tailed confidence interval for difference of means: 83.694 to 113.056

Two-tailed P-value = 2.262E-012

**t-test**

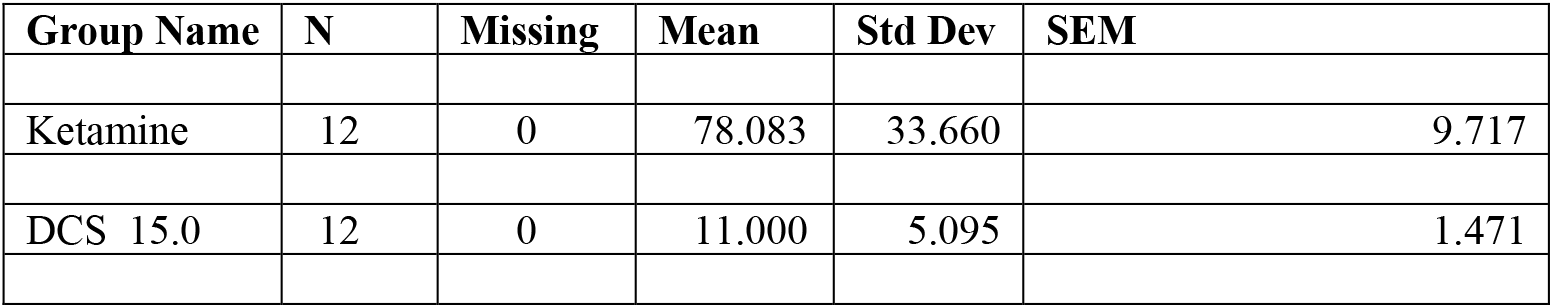

Difference 67.083 t = 6.826 with 22 degrees of freedom.

95 percent two-tailed confidence interval for difference of means: 46.702 to 87.464

Two-tailed P-value = 0.000000740

